# Cotranscriptional kinetic folding of RNA secondary structures

**DOI:** 10.1101/2020.07.10.196972

**Authors:** Vo Hong Thanh, Pekka Orponen

**Affiliations:** Department of Computer Science, Aalto University

**Keywords:** RNA secondary structure, Costranscriptional folding, Stochastic simulation

## Abstract

Computational prediction of RNA structures is an important problem in computational structural biology. Studies of RNA structure formation often assume that the process starts from a fully synthesized sequence. Experimental evidence, however, has shown that RNA folds concurrently with its elongation. We investigate RNA structure formation, taking into account also the cotranscriptional effects. We propose a single-nucleotide resolution kinetic model of the folding process of RNA molecules, where the polymerase-driven elongation of an RNA strand by a new nucleotide is included as a primitive operation, together with a stochastic simulation method that implements this folding concurrently with the transcriptional synthesis. Numerical case studies show that our cotranscriptional RNA folding model can predict the formation of metastable conformations that are favored in actual biological systems. Our new computational tool can thus provide quantitative predictions and offer useful insights into the kinetics of RNA folding.

## 1 Introduction

Ribonucleic acid (RNA) is a biopolymer constituted of nucleotides with bases adenine (A), cytosine (C), guanine (G) and uracil (U). The synthesis of an RNA molecule from its DNA template is initiated when the corresponding RNA polymerase binds to the DNA promoter region. RNA has been shown to serve diverse functions in a wide range of cellular processes such as regulating gene expression and acting as an enzymatic catalyst [3, 23], and has also recently been used as an emerging material for nanotechnology [11].

Computational prediction of RNA secondary structures given their primary sequences is often based on the estimation of changes in free energy, which postulates that thermodynamically an RNA strand will fold into a conformation that yields the minimum free energy (MFE) value (see e.g., [5] for a review on the topic). We limit our focus in this work to *pseudoknot-free* secondary structures, i.e., structures with no crossing base pairs, because finding MFE structures with pseudoknots in a general energy model is an NP-complete problem [15]. Under this simplification, the energy of an RNA secondary structure can be modeled as the sum of energies of strand loops flanked by base pairs. The loop energy parameters have been measured experimentally and are detailed in a nearest neighbor parameter database [27]. Methods grounded in the thermodynamic framework, such as the Zuker algorithm [31] and its extensions [30, 17], can be used to compute pseudoknot-free MFE secondary structures effectively in a bottom-up manner.

The kinetic approach is an alternative way to study the RNA folding process. It models the folding as a random process where the additions/deletions of base pairs in the current structure are assigned probabilities proportional to the respective changes in free energy values [6]. A folding pathway of a sequence is then generated by executing a stochastic simulation computation [6, 19, 4]. We refer to the book by Marchetti *et al.* [16] for a comprehensive review of state-of-the-art stochastic simulation techniques. Each simulation run on a given RNA sequence can produce a list of possible structures that it can fold into. Such dynamic view of RNA folding allows one to capture cases where local conformations are progressively folded to create metastable structures that kinetically trap the folding, thus complementing the prediction of equilibrium MFE structures produced by the thermodynamic approach.

The study of RNA structure formation often assumes that the folding process starts from a fully synthesized open strand, the *denatured state*. However, experimental evidence [28, 21] has shown that RNA starts folding already concurrently with the transcription. The nucleotide transcription speed varies from 200 nt/sec (nucleotides per second) in phages, to 20-80 nt/sec in bacteria, and 5-20 nt/sec in humans [21]. The RNA dynamics also occur over a wide range of time-scales where base pairing takes about 10^−3^ msec; structure formation is about 10-100 msec; and kinetically trapped conformations can persist for minutes or hours [28]. One consequence of considering cotranscriptional folding is that the base pairs at the 5’ end of the RNA strand will form first, while the ones at the 3’ end can only be formed once the transcription is complete, which leads to structural asymmetries. Cotranscriptional folding can thus form *transient* structures that are only present for a specific time period and involved in distinct roles. For instance, gene expression when considering such transient conformations of RNA during cotranscriptional folding can expose oscillation behavior [2]. We refer to the review by Lai *et al.* [14] for further discussion on the importance of cotranscriptional effects.

In this work, we extend the kinetic approach to take into account cotranscriptional effects on the folding of RNA at single-nucleotide resolution. Unlike existing kinetic methods that either start from a fully synthesized sequence [6] or approximate the effects of co-transcriptional folding [22, 20, 10, 29, 9], we explicitly consider the elongation of RNA during transcription as a primitive action in the model. The time when a new nucleotide is added to the current RNA chain is specified by the transcription speed of the RNA polymerase enzyme. The RNA strand in our modeling approach can elongate with newly synthesized nucleotides added to the sequence and fold simultaneously. We also propose an exact stochastic simulation method, the CoStochFold algorithm, to handle the transcription events.

The rest of the paper is organized as follows. Sec. 2 reviews some background on kinetic folding of RNA. In Sec. 3, we present our work to extend the model of RNA folding to incorporate the transcription process. Sec. 4 reports our numerical experiments on case studies. Concluding remarks are in Sec. 5.

## 2 Background on kinetic folding

Let *S_n_* be a linear sequence of length *n* of four bases A, C, G, and U in which the 5’ end is at position 1 and the 3’ end is at position *n*. A base at position *i* may form a hydrogen bond with a base at *j*, denoted by (*i, j*), if they form a *Watson-Crick pair* A-U, G-C or a *wobble pair* G-U. A secondary structure formed by intra-molecular interactions between bases in *S_n_* is a list of base pairs (*i, j*) with *i* < *j* satisfying constraints: a) the *i*th base and *j*th base must be separated by at least 3 (un-paired) bases, i.e., *j* − *i* > 3; b) for any base pair (*k, l*) with *k* < *l*, if *i* = *k* then *j* = *l*; and c) for any base pair (*k, l*) with *k* < *l*, if *i* < *k* then *i* < *k* < *l* < *j*. The first condition prevents the RNA backbone from bending too sharply. The second one prevents the forming of tertiary structure motifs such as base triplets and G-quartets. The last constraint ensures that no two base pairs intersect, i.e. there are no pseudoknots.

Let 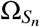 be the set of all possible secondary structures formed by *S_n_*. Consider a secondary structure 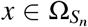. It can be represented compactly as a string of dots and brackets (see Fig. 1). Specifically, for a base pair (*i, j*), an opening bracket ‘(’ is put at *i*th position and a closing bracket ‘)’ at *j*th position. Finally, unpaired positions are represented by dots ‘.’. The dot-bracket representation is unambiguous because the base pairs in a secondary structure do not cross each other.

**Fig. 1.**
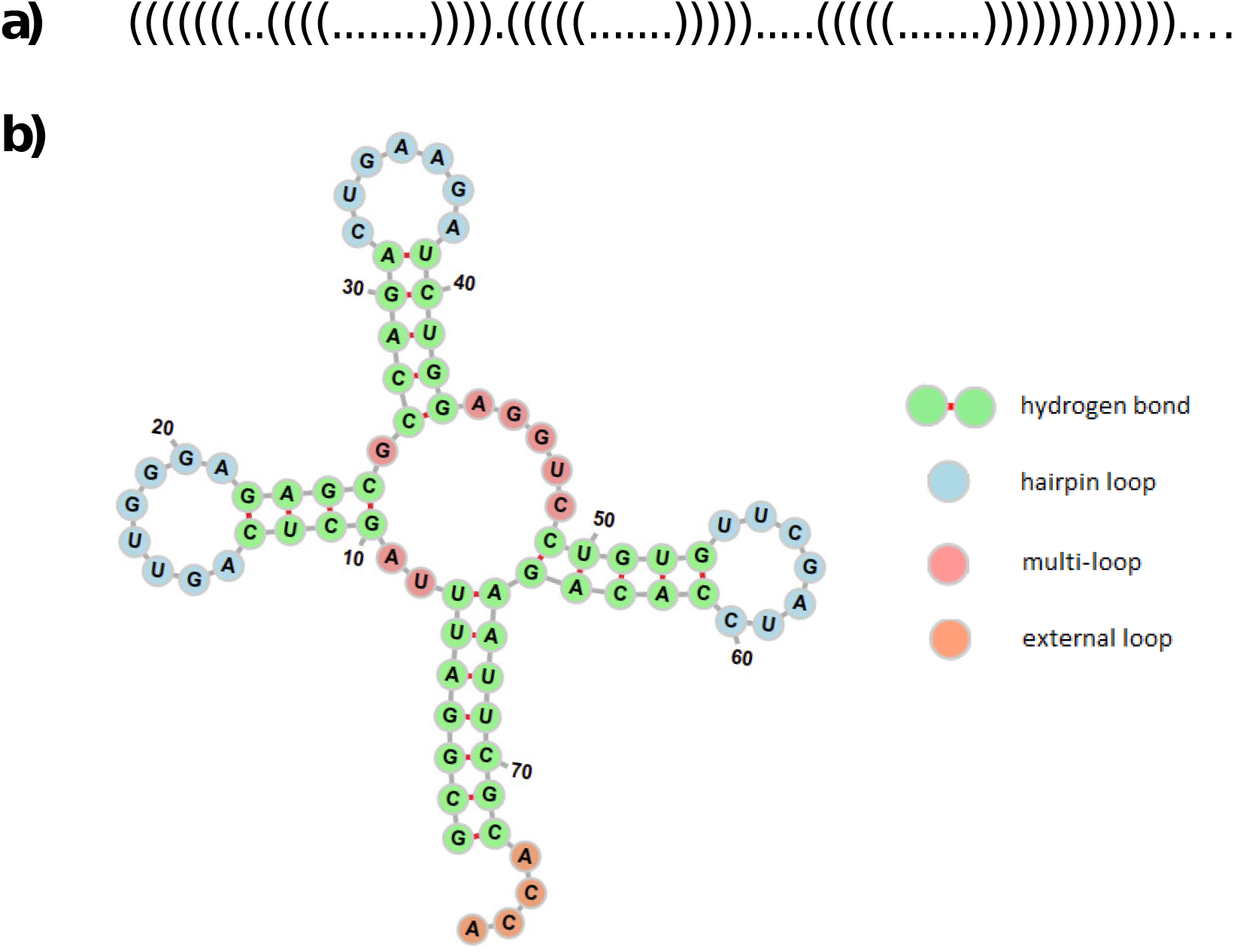
Representation of the tRNA molecule in a) dot-bracket notation and b) graphical visualization. The graphical visualization is made by the Forna tool [13].

The free energy of *x* can be estimated by the *nearest neighbor* model [17], in which the free energy of an RNA secondary structure is taken to be the sum of energies of components flanked by base pairs. Formally, for a base pair (*i, j*) in *x*, we say that base *k*, *i* < *k* < *j*, is *accessible* from (*i, j*) if there is no other base pair (*i′, j′*) such that *i* < *i′* < *k* < *j′* < *j*. The set of accessible bases flanked by base pair (*i, j*) is called the *loop* **L**(*i, j*). The number of unpaired bases in a loop **L**(*i, j*) is its *size*, while the number of enclosed base pairs determines its degree. Based on these properties, loops **L**(*i, j*) can be classified as *stacks*, *hairpins*, *bulges*, *internal loops* and *multi-loops*. The unpaired bases that are not contained in loops constitute the *exterior* (or external) loop **L**_*e*_.

A secondary structure *x* is thus uniquely decomposed into a collection of loops *x* = ⋃_(*i, j*)_**L**(*i, j*)⋃**L**_*e*_. Based on this decomposition, the free energy *G_x_* (in kcal) of secondary structure *x* is computed as:

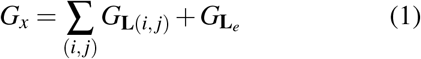

where *G*_**L**(*i, j*)_ is the free energy of loop **L**(*i, j*). Experimental energy values for *G*_**L**(*i, j*)_ are available in the nearest neighbor database [27].

Let 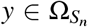 be a secondary structure derived directly from *x* by an intramolecular reaction between bases *i* and *j* in *x*. Commonly, three operations on a pair of bases, referred to as the *move set* (see Fig. 2), are defined [6]:

*Addition: y* is derived by adding a base pair that joins bases *i* and *j* in *x* that are currently unpaired and eligible to pair.
*Deletion: y* is derived by breaking a current base pair (*i, j*) in *x*.
*Shifting: y* is derived by shifting a base pair (*i, j*) in *x* to form a new base pair (*i, k*) or (*k, j*).

**Fig. 2.**
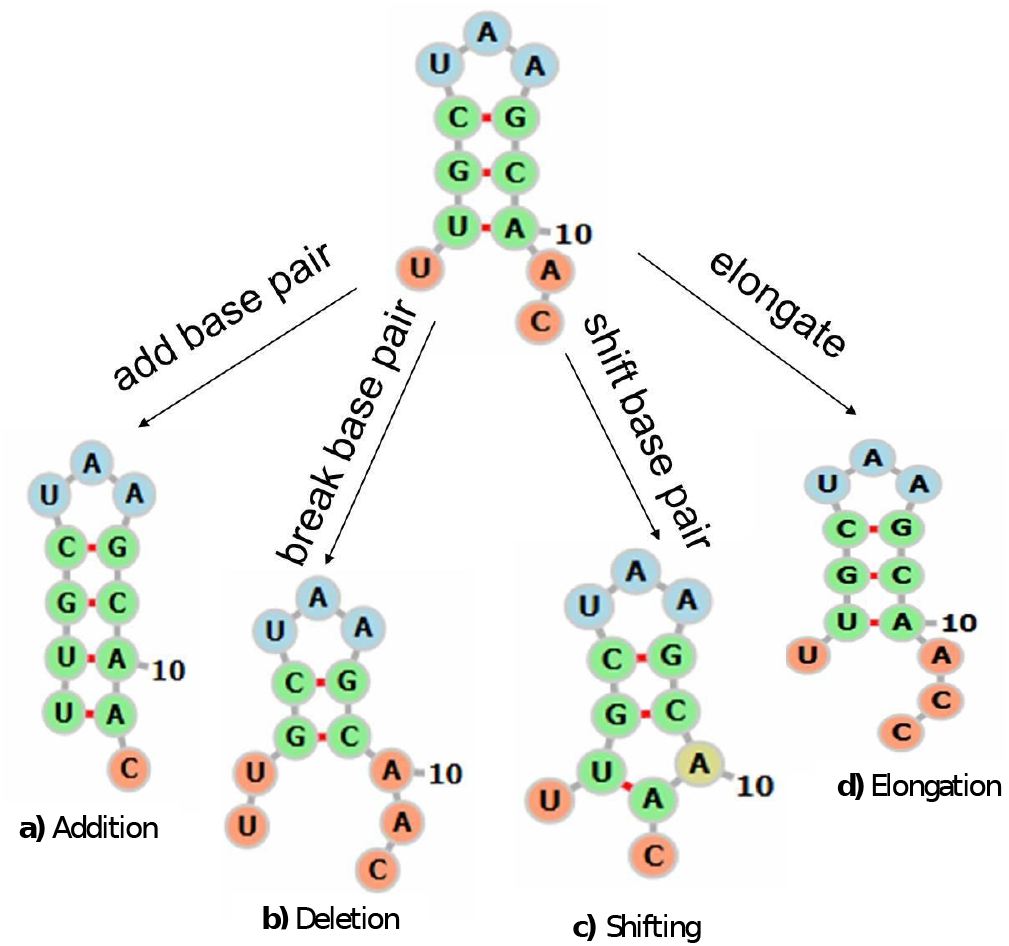
Extended move set consisting of a) addition, b) deletion, c) shifting and d) elongation. The elongation move models the transcription process extending the current RNA chain with a new nucleotide at the 3’ end.

Let *k_x→y_* be the rate (probability per time unit) of the transition from *x* to *y*. In a conformation *x*, the RNA molecule may wander vibrationally around its energy basin *G_x_* for a long time, before it surmounts an energy barrier to escape to a conformation *y* in another basin. The dynamics of the transition from *x* to *y* characterizes a rare event in Molecular Dynamics (MD). Here, we adopt the coarse-grained kinetic Monte Carlo approximation [18, 12], and model the transition rate *k_x→y_* as:

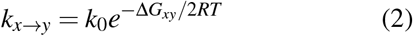

where *T* is absolute temperature in Kelvin (K), *R* = 1.98717 × 10^−3^(*kcal K^−^*^1^ · *mol^−^*^1^) is the gas constant and Δ*G_xy_* = *G_y_* − *G_x_* denotes the difference between free energies of *x* and *y*. *k*_0_ is a constant for calibration of time.

Let *P*(*x,t*) be the probability that the system is at conformation *x* at time *t*. The dynamics of *P*(*x,t*) is formulated by the (chemical) master equation [16] as:

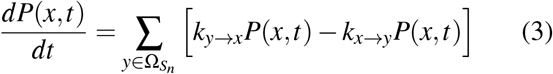

Analytically solving Eq.3 requires to enumerate all possible states *x* and their neighbors *y*. The size of the state space 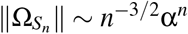 with α = 1.8488 increases exponentially with the sequence length *n*, and the number of neighbors of *x* is in order of *O*(*n*^2^) [8]. Thus, due to the high dimension of the state space, solving Eq.3 often involves numerical simulation.

Let *P*(*y*, τ|*x,t*) be the probability that, given current structure *x* at time *t*, *x* will fold into *y* in the next infinitesimal time interval [*t* + τ,*t* + τ+ *d*τ). We have

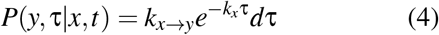

where 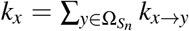 is the sum of transition rates to single-move neighbors of *x*. Eq. 4 shows that the probability that *x* moves to *y* is *k_x→y_*/*k_x_*, and the waiting time τ until that transition follows an exponential distribution *Exp*(*k_x_*). These facts are the basis for our kinetic folding algorithm *StochFold* presented as Algorithm 1. We note that StochFold shares the structure of the earlier algorithm Kinfold [6] and its improvements [4, 25, 24, 26].

**Algorithm 1.**
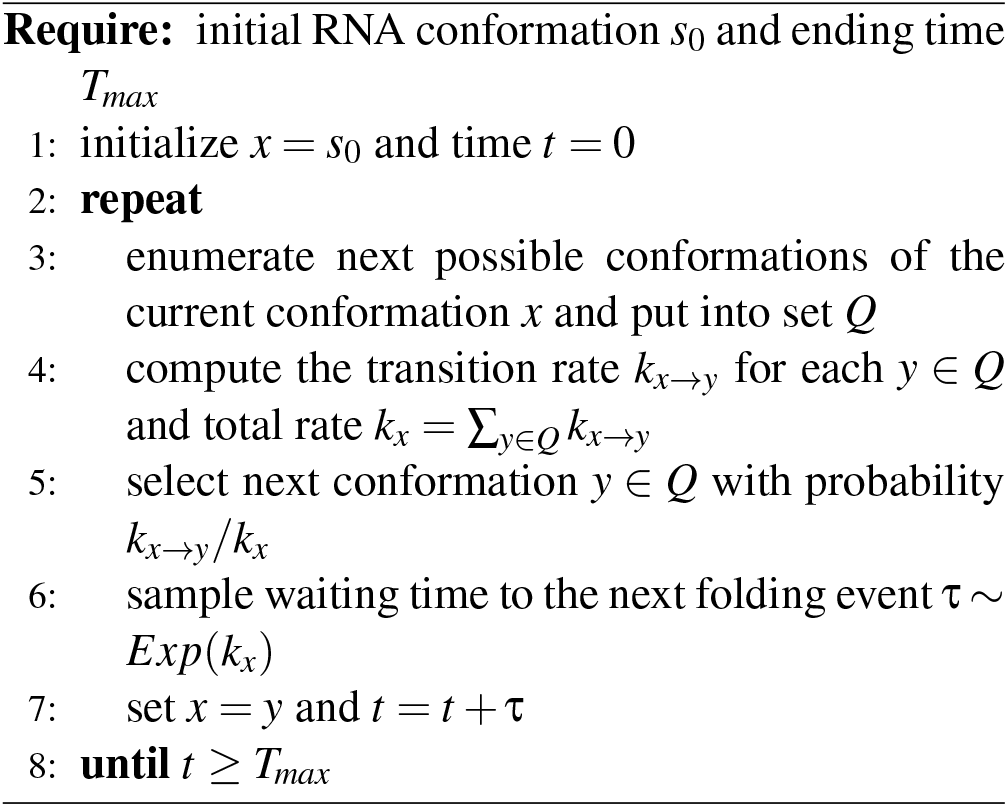
StochFold

## 3 Cotranscriptional kinetic folding of RNA

The folding of an RNA strand adapts immediately to new nucleotides synthesized during the transcription. The kinetic approach described in Section 2 cannot capture the effects of such cotranscriptional folding, because it considers only interactions between bases already present in the sequence. We outline in this section an approach to incorporating these effects in the simulation. The transcription process is explicitly taken into account by extending the move set with the new operation of *elongation*. Our extended move set thus comprises four operations: addition, deletion, shifting and elongation. The first three operations are defined as in the previous section. In elongation, the current RNA chain increases in length and a newly synthesized nucleotide is added to its 3’ end. Figure 2 illustrates the extended move set.

Under the extended move set, we define two event types: *folding event* and *transcription event*. A folding event is an *internal event* that occurs when one of the three operations addition, deletion or shifting, is applied to a base pair of the current sequence. A transcription event happens when the elongation operation is applied. It is an *external event* whose rate is specified by the transcription speed of the RNA polymerase enzyme. The occurrences of transcription events break the Markovian property of transitions between conformations. This is because when a new nucleotide is added to the current RNA conformation, the number of next possible conformations increases. The waiting time of the next folding event also changes and thus a new folding event has to be recomputed.

Algorithm 2 outlines how the CoStochFold algorithm handles this situation. The key element of CoStochFold (lines 8 - 14) is a race where the event having the smallest waiting time will be selected to update the current RNA conformation. More specifically, suppose the current structure is *x* at time *t*. Let τ_*e*_ be the waiting time to the next folding event and τ_*trans*_ the waiting time to the next transcription event. Assuming that no events occur earlier, τ_*e*_ has an exponential distribution with rate *k_x_* which is the sum of all transition rates of applying addition, deletion and shifting operations to base pairs in *x*. For simplifying the computation of τ_*trans*_, we assume that it is the expected time to transcribe one nucleotide. Let *N_trans_* be the (average) transcription speed of the polymerase. We compute τ_*trans*_ as:

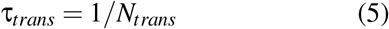

Thus, given current time *t*, the next folding event will occur at time *t_e_* = *t* + τ_*e*_ and, respectively, the transcription event where a new nucleotide will be added to the current sequence is scheduled at time *t_trans_* = *t* + τ_*trans*_. We decide which event will occur by comparing *t_e_* and *t_trans_*. If *t_e_* > *t_trans_*, then a new nucleotide is first transcribed and added to the current RNA conformation. Otherwise, a folding event is performed where a structure in the set *Q* of neighboring structures is selected to update the current conformation.

We remark that one can easily extend Algorithm 2 to allow modeling τ_*trans*_ as a random variable without changing the steps of event selection. Specifically, one only needs to change step 2 in Algorithm 2 to generate the waiting time of the next transcription event, while keeping the simulation otherwise unchanged.

## 4 Numerical experiments

We illustrate the capability of our cotranscriptional kinetic folding method on three case studies: a) the switching molecule [6], b) the E. coli signal recognition particle (SRP) RNA and c) the SV-11 variant in Qβ replicase [1]. SRP and SV-11 are real biological molecules that manifest the characteristics of cotranscriptional folding that thermodynamic/kinetic methods [31, 7] would fail to capture if initiated from fully denatured sequences. Our method is not only able to produce these structures, but also provides insight into mechanisms that biological systems may use to guide the structure formation process. The code for the implementation of our CoStochFold algorithm is available at: https://github.com/vo-hong-thanh/stochfold.

**Algorithm 2.**
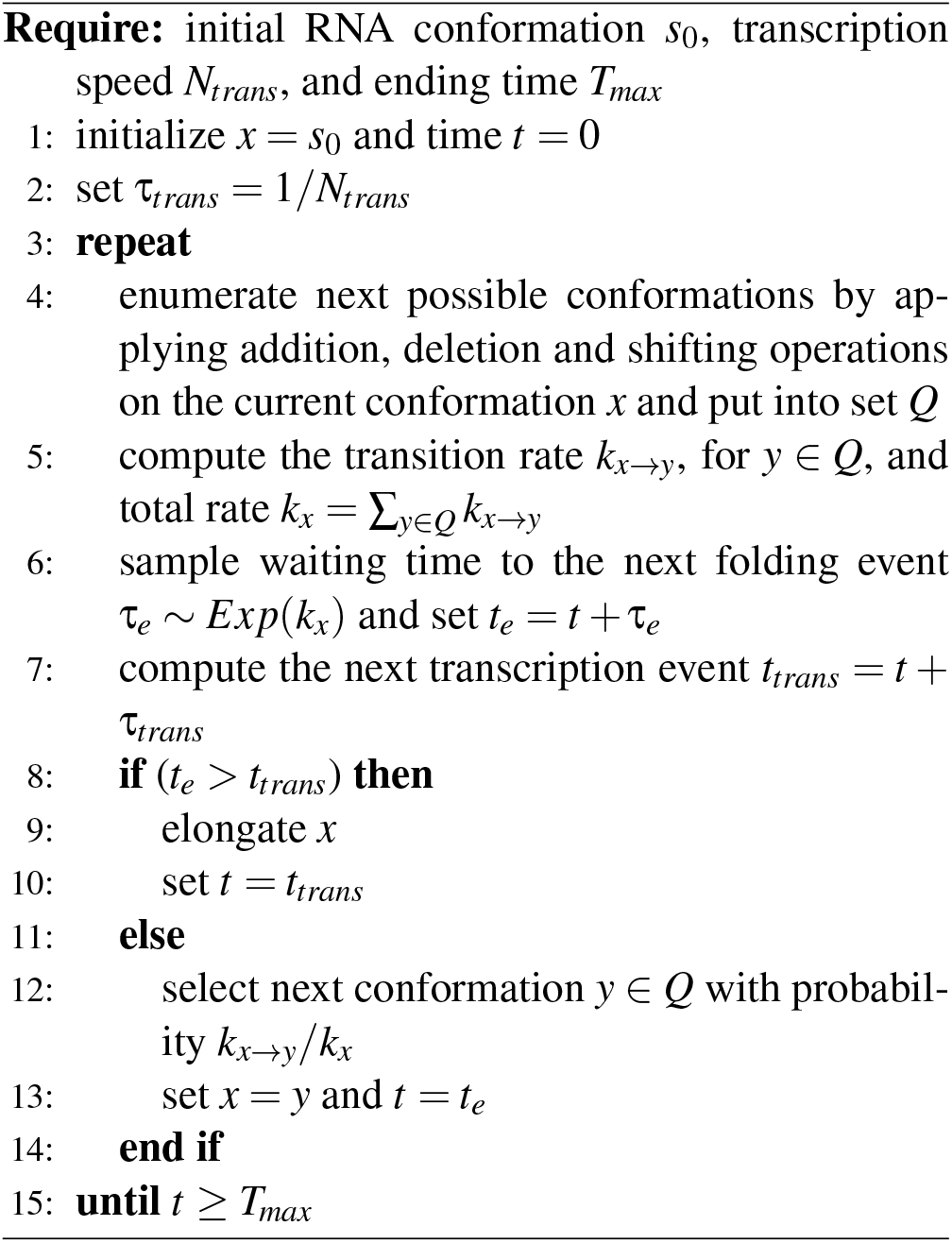
CoStochFold

### 4.1 Switching molecule

We consider the dynamic folding of the RNA sequence *S* = “GGCCCCUUUGGGGGCCAGACC-CCUAAAGGGGUC” [6]. Two stable conformations of the sequence are: the MFE structure *x* = “((((((((((((((.….))))))))))))))” (−26.20 kcal), and a suboptimal structure *y* = “((((((.…)))))).((((((.))))))” (−25.30 kcal). In this example, we focus on the number of first-hitting time occurrences of a target structure. The number of first-hitting time occurrences of a structure in a time interval divided by the total number of simulation runs approximates the first-passage time probability of the structure, i.e., its folding time [6].

Fig. 3 plots the number of first-hitting time occurrences of the MFE structure *x* and the suboptimal *y* with varying transcription speeds. Kinetic folding starting from the denatured state was carried out by the StochFold algorithm, while cotranscriptional folding was conducted by the CoStochFold algorithm. For each method, we performed 10000 simulation runs on the sequence *S* in which each simulation ran until a target structure was observed or the ending time *T_max_* = 1000 was reached. Fig. 3 shows that changing the transcription speed of the polymerase significantly affects the folding characteristics of the sequence. Specifically, co-transcriptional folding with slow transcription speed favors the suboptimal structure *y*. It increases the number of occurrences of *y*, while reducing the number of occurrences of the MFE structure *x*.

**Fig. 3.**
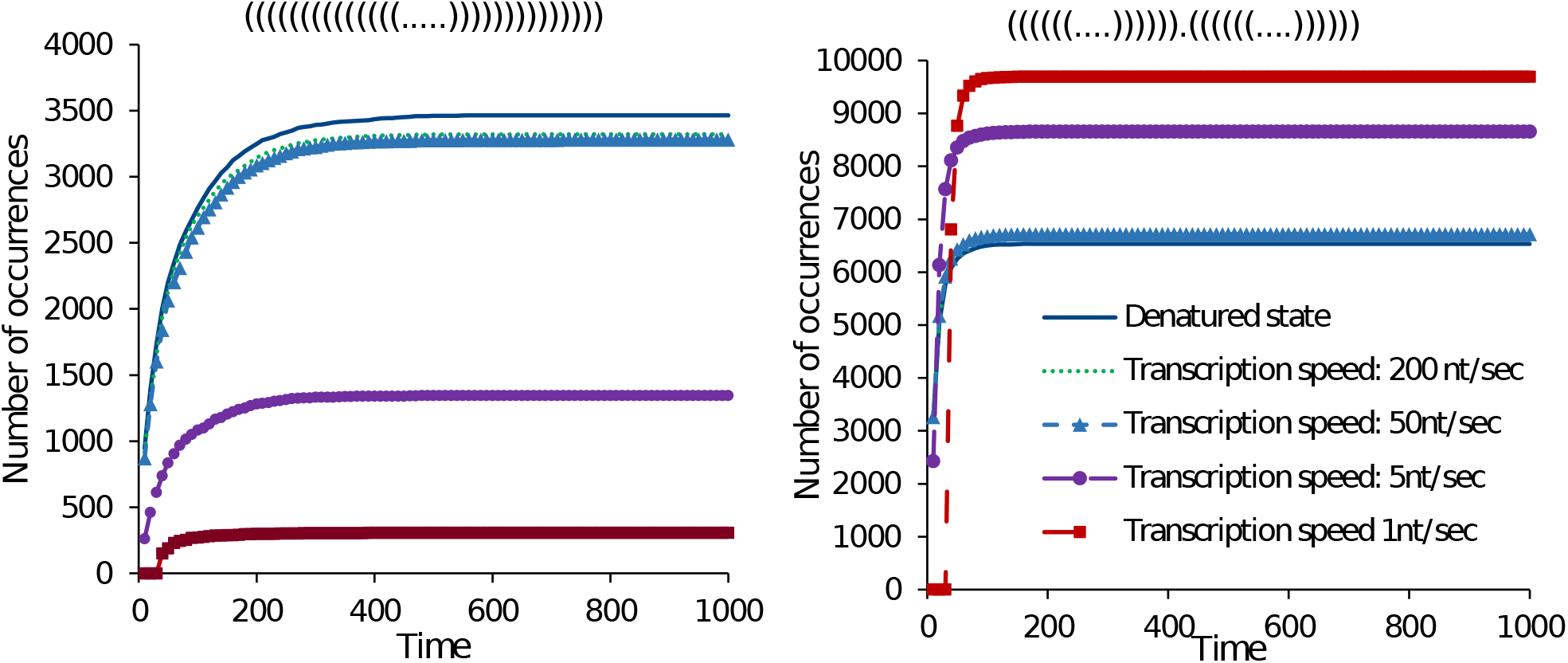
First-hitting time occurrences of MFE structure *x* = “((((((((((((((.….))))))))))))))” (−26.20 kcal, left) and suboptimal *y* = “((((((.…)))))).((((((.))))))” (−25.30 kcal, right).

Fig. 4 compares the number of first-hitting time occurrences of the MFE structure *x* with respect to the suboptimal conformation *y*. We note that if the simulation starts from the fully denatured state, the occurrence ratio of the suboptimal conformation *y* to the MFE structure *x* is about 2:1 as also observed by Flamm *et al.* [6]. However, the ratio increases noticeably when the transcription speed decreases. For example, the occurrence ratio of the suboptimal conformation *y* to the MFE structure *x* is about 6.5:1 in the case of transcription speed 5 nt/sec.

**Fig. 4.**
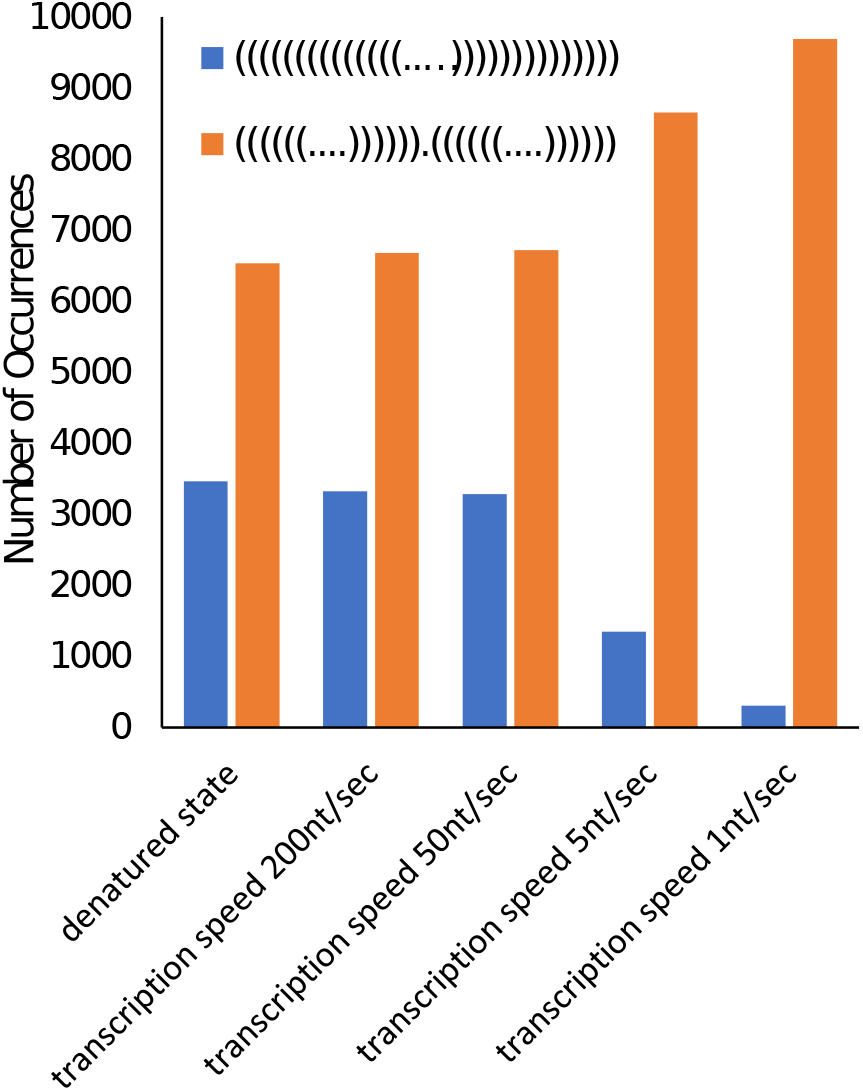
First-hitting time occurrences of MFE structure *x* = “((((((((((((((.….))))))))))))))” (−26.20 kcal) and suboptimal *y* = “((((((.…)))))).((((((.…))))))” (−25.30 kcal) by varying transcription speeds.

### 4.2 Signal recognition particle (SRP) RNA

This section studies the process of structural formation of the E. coli SRP RNA during transcription. SRP is a 117nt long molecule, which recognizes the signal peptide and binds to the ribosome locking the protein synthesis. Its active structure is a long helical structure containing interspersed inner loops (see S3 in Fig. 5). Experimental work [28] using SHAPE-seq techniques has suggested a series of structural rearrangements during transcription that ultimately result in the SRP helical structure. In particular, the 5’ end of SRP forms a hairpin structure during early transcription. The structure persists until it reaches a length of 117nt. The unstable hairpin then rearranges to its active structure. Fig. 5 depicts three structural motifs at 25nt (S1), 86nt (S2), and 117nt (S3), respectively, in the formation of SRP. Specifically, the hairpin motif S1 emerges at transcript length 25nt, and the transcript then continues elongating to form structure S2 at length 86nt. When reaching the transcript length 117nt, SRP rearranges into its persistent helical conformation S3.

**Fig. 5.**
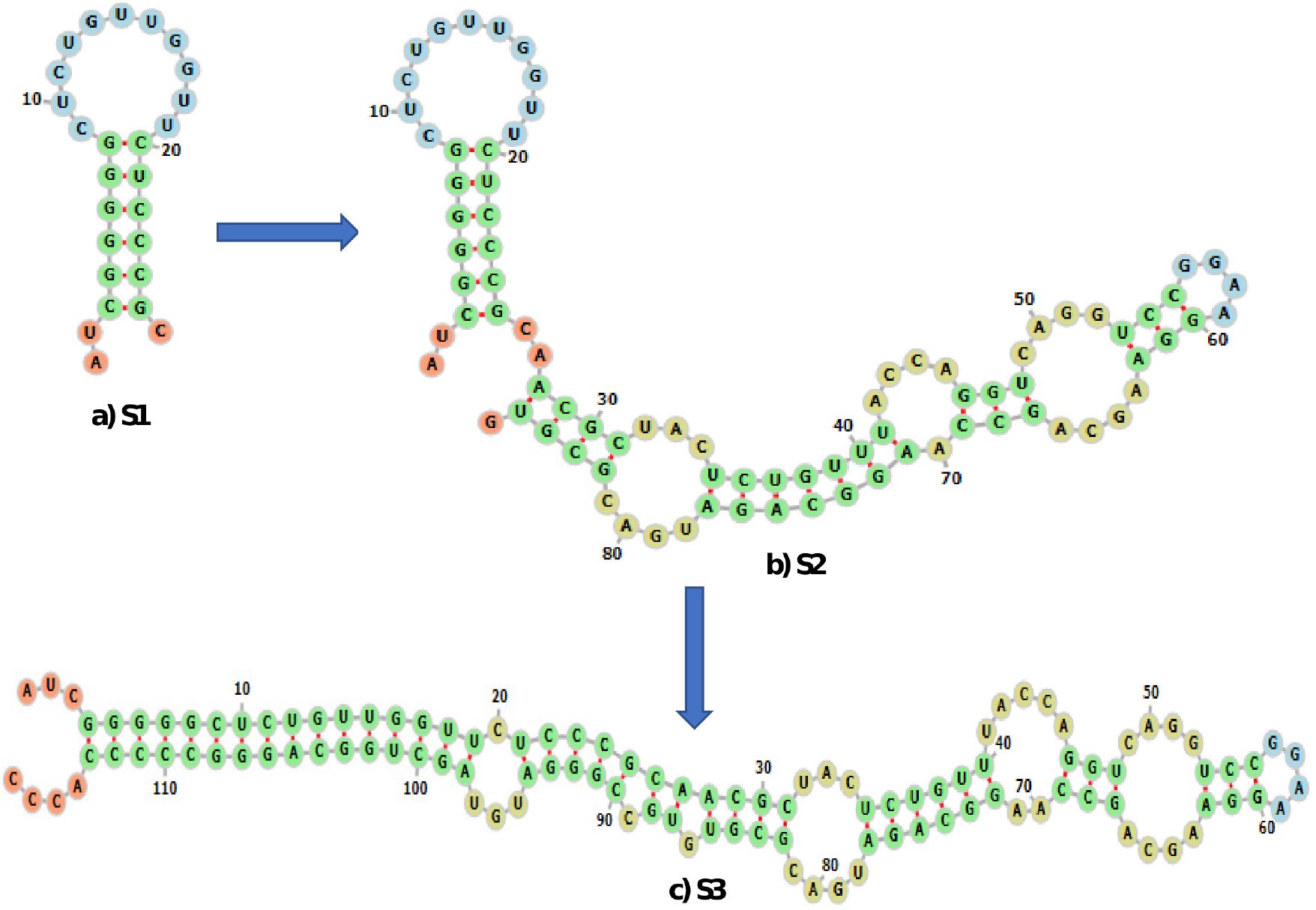
The folding pathway of secondary structures of E. Coli signaling recognition particle (SNP) RNA. The hairpin motif S1 is formed at transcript length 25nt and form S2 completed at length 86nt. When reaching the transcript length 117nt, SRP rearranges into its stable helical shape S3. The visualization of structures is made by the Forna tool [13].

We validated the prediction of the CoStochFold algorithm against the experimental work in [28]. To do that, we performed 10000 simulations of the algorithm to fold SRP transcriptionally. The average transcription speed was set to 5 nt/sec. Fig. 6 shows the frequency of occurrences of the considered structures during the simulated time of 30 seconds. The plot on the left shows the cotranscriptional folding of SRP and the plot on the right presents the folding of SRP starting from the denatured state. The figures clearly show that the CoStochFold algorithm can capture the folding pathway of SRP. Specifically, the hairpin motif S1 starts to form at about *t* = 4s when the transcript length is 20nt and peaks at about *t* = 8s when 40nt have been transcribed. At about *t* = 18s, Structure S2 appears and then rearranges to S3 at about *t* = 24s. We note that the simulated folding without considering transcription only the conformation S3 is found.

**Fig. 6.**
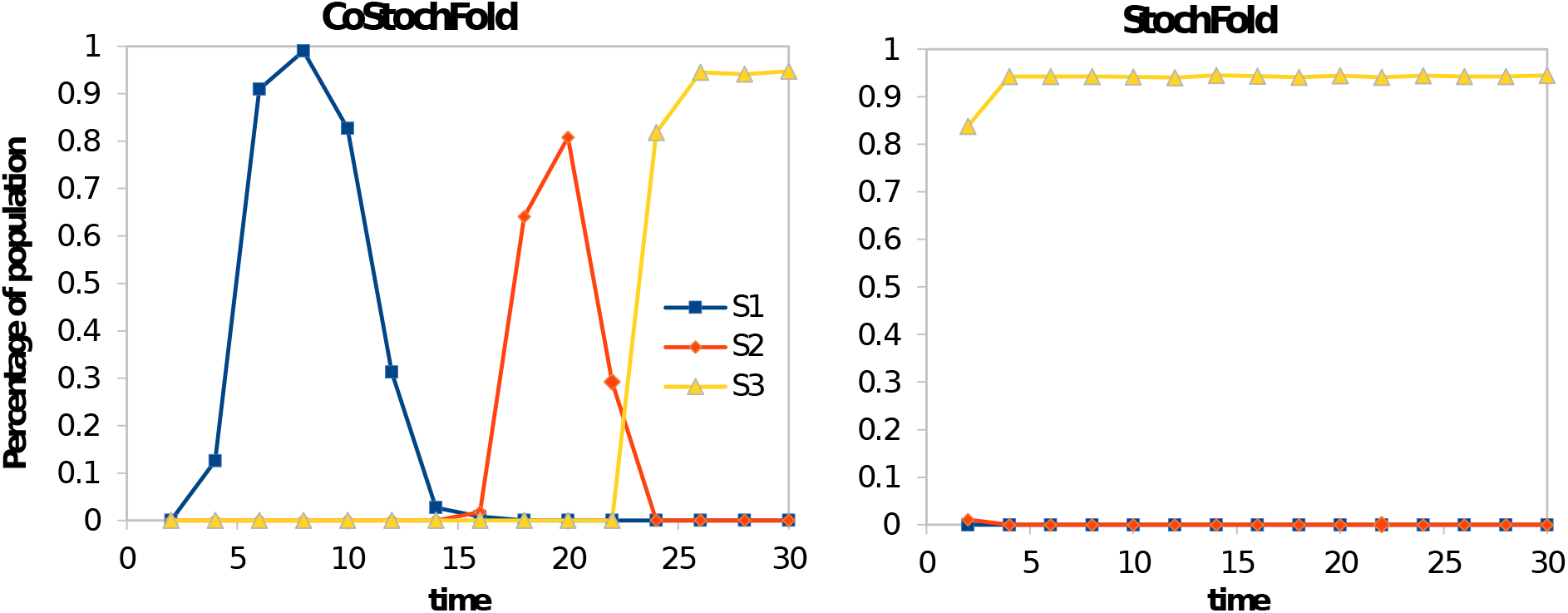
Prediction of the structural formation of SNP. Left: cotranscriptional folding. Right: folding from denatured state without transcription.

### 4.3 SV-11

SV-11 is a 115 nt long RNA sequence. It is a recombinant between the plus and minus strands of the natural Qβ template MNV-11 RNA [1]. The result of the recombination is a highly palindromic sequence whose most stable secondary structure is a long hairpin-like structure, the MFE structure in Fig. 7a). The MFE structure, however, disables Qβ replicase because its primer regions are blocked. Experimental work [1] has shown that an active structure of SV-11 for replication is when it folds into a metastable conformation. Fig. 7b) depicts a metastable structure with a hairpin-hairpin-multi-loop motif with open primer regions that serve as template for replication. Transition from the metastable structure to the MFE structure has been observed experimentally but is rather slow [1], indicating long relaxation time to equilibrium.

**Fig. 7.**
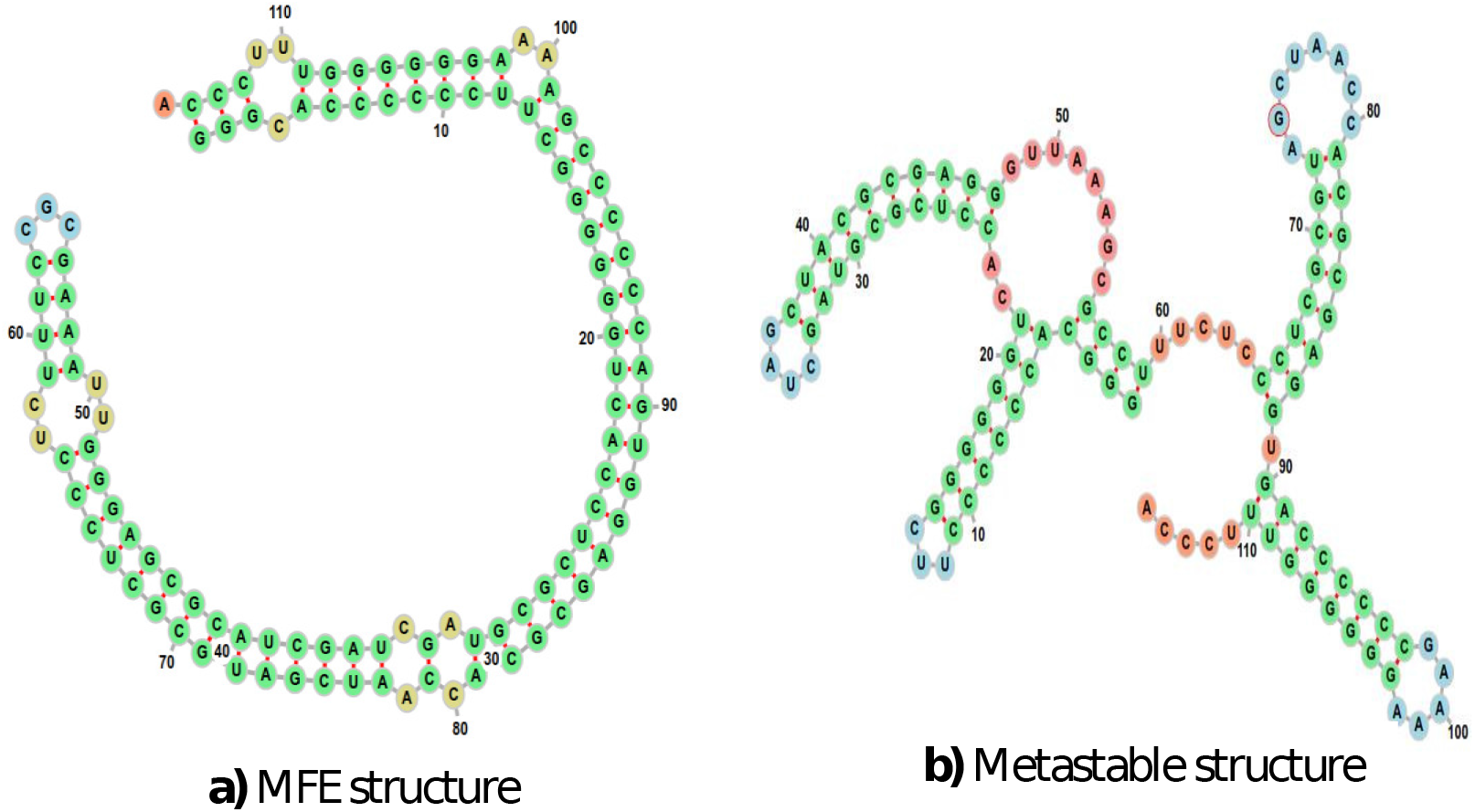
SV-11 with two conformations a) MFE structure (−95.90 kcal) and b) metastable structure (−63.60 kcal). The visualization of structures is made by the Forna tool [13].

We plot in Fig. 8 the energy vs. occurrence frequency of structures by the cotranscriptional folding of SV-11. The result is obtained by 10000 simulation runs of our CoStochFold algorithm with average transcription speed 5nt/sec. To determine the frequency of occurrence of a structure, we discretize the simulation time [0, *t*] into intervals and record how much time was spent in each structure within each interval. The frequency of occurrence of a structure in each time interval is then averaged over 10000 runs. The figure shows that the folding favors metastable structures, and disfavors the MFE structure. In particular, cotranscriptional folding quickly folds SV-11 to its metastable conformations with the mode of the energy distribution at about −63kcal.

**Fig. 8.**
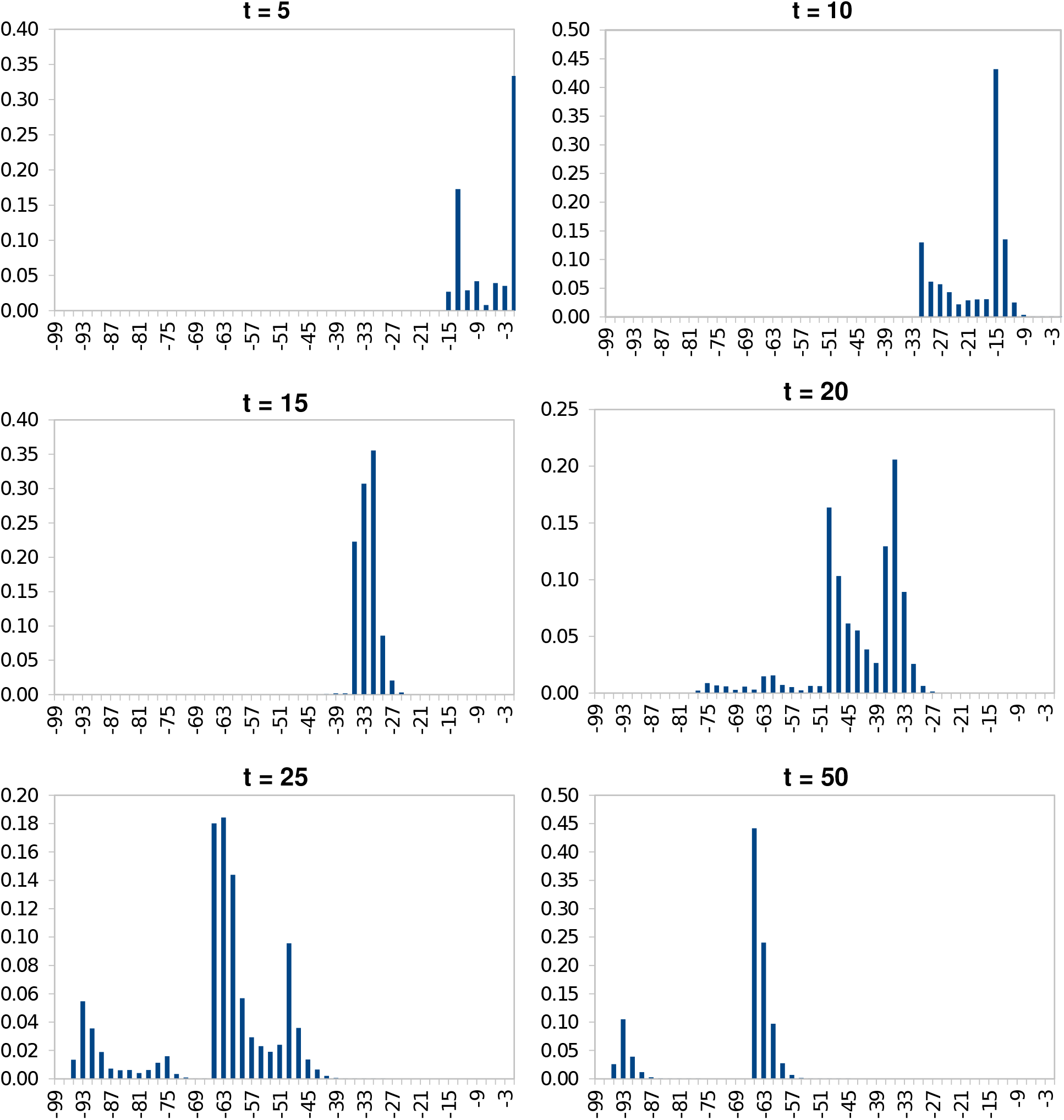
Cotranscriptional folding of SV-11. The x-axis denotes the energy level, and y-axis shows the frequency of structures in the same energy level.

Fig. 9 shows the long-term occurrence frequencies of structures at different energy levels in the SV-11 folding and Fig. 10 compares the occurrence frequencies of a specific metastable structure depicted in Fig. 7b) with the MFE structure and two randomly selected suboptimal structures in the locality of MFE structure. Fig. 10 shows that the SV-11 molecule interestingly prefers the metastable structure over the MFE structure. Specifically, the metastable structure in the cotranscriptional folding regime is in the time interval [0, 10000] about ten-fold more frequent than the MFE structure.

**Fig. 9.**
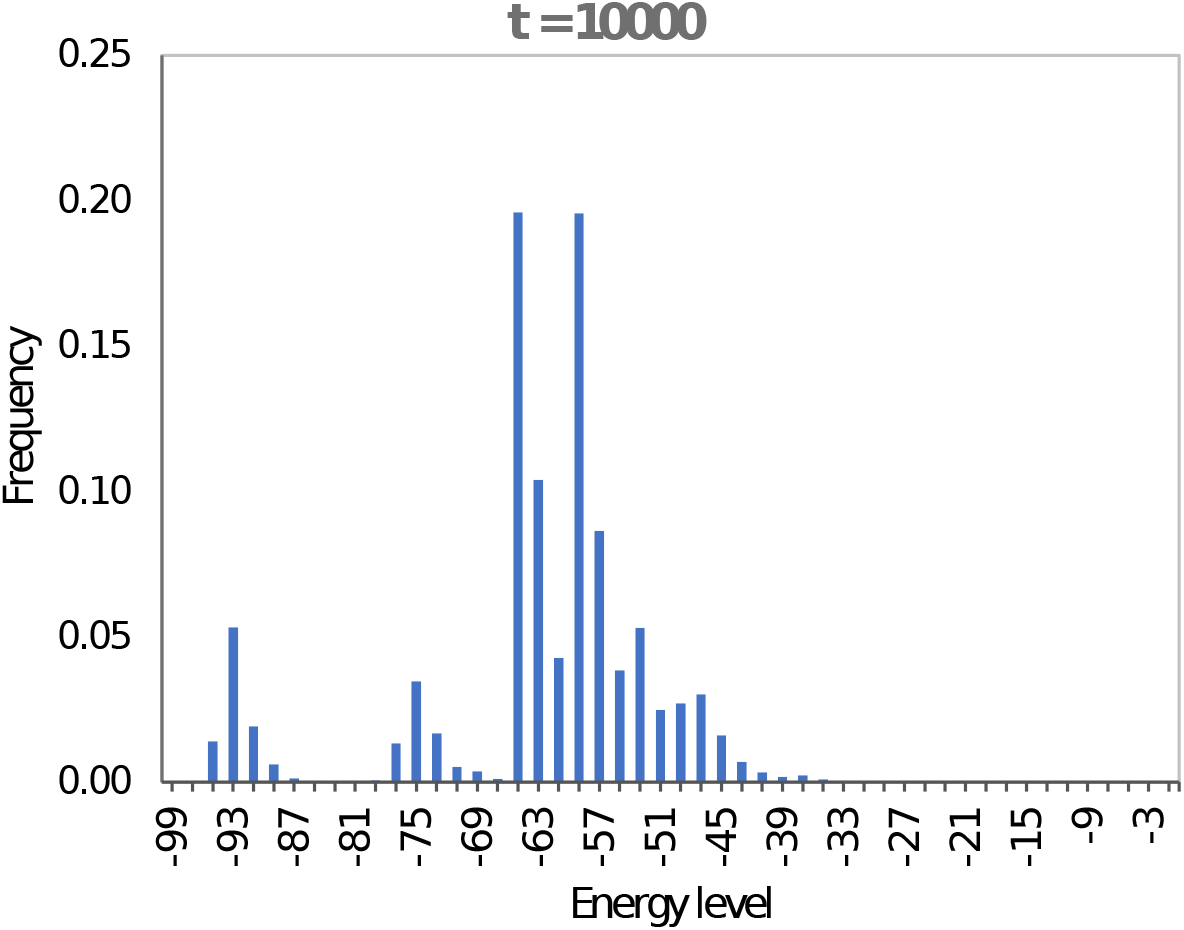
Frequency of structures in folding SV-11.

**Fig. 10.**
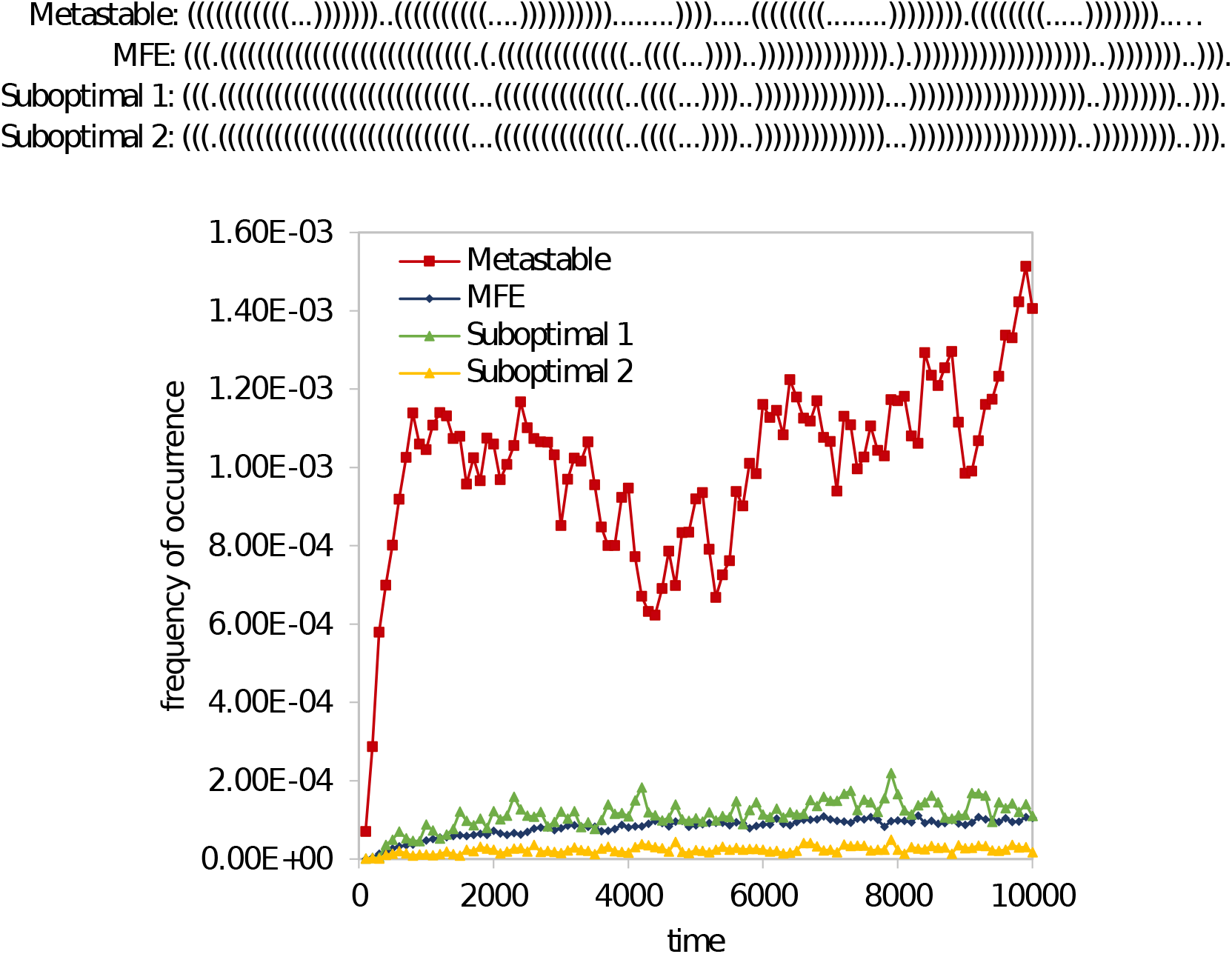
Frequency of the metastable structure in comparison with the MFE structure and two randomly selected suboptimal structures.

## 5 Conclusions

We propose a kinetic model of RNA folding that takes into account the elongation of an RNA chain during transcription as a primitive structure-forming operation alongside the common base-pairing operations. Based on this, we developed a new stochastic simulation algorithm CoStochFold to explore RNA structure formation in the cotranscriptional folding regime. We showed through numerical case studies that the method can quantitatively predict the formation of metastable conformations in an RNA folding pathway. The simulation method and tools thus promise to offer useful insights into RNA folding kinetics in real biological systems.

## Acknowledgements

This work has been supported by Academy of Finland grant no. 311639, “Algorithmic Designs for Biomolecular Nanostructures (ALBION).”

